# Constant companions: Wild zebra finch pairs display extreme spatial cohesion

**DOI:** 10.1101/2024.03.21.586046

**Authors:** Chris Tyson, Hugo Loning, Simon C. Griffith, Marc Naguib

## Abstract

Many animals maintain long-term monogamous partnerships, but the extent to which partners associate varies substantially and has implications for the scope of cooperation between pair members. Zebra finches (*Taeniopygia castanosis*) are monogamously paired for life and maintain continuous partnerships, raising questions as to if and how they maintain pair cohesion despite being nonterritorial and having only short-range acoustic signals. While zebra finches are the best studied songbird in captivity, their social and spatial behaviour in the wild is poorly understood. Determining pair cohesion in songbirds to date has almost exclusively been studied at specific locations where pairs would be expected to meet, such as nesting or feeding sites, without quantifying broader movements. Here, we used solar-powered automated tracking to simultaneously monitor the movements of radio-tagged zebra finch pairs during periods with breeding activity. We reveal extremely high spatial cohesion with pairs using nearly identical home ranges and maintaining close spatial proximity across large areas. This characterisation of extremely high spatio-temporal coordination of zebra finch pairs provides important insights into the operation and benefits of monogamous relationships in highly mobile taxa, such as birds.

## I. Introduction

Socially monogamous relationships in animals have been characterised by surprisingly long associations between males and females that can last over decades in birds (e.g., [1,2]) or as long as 27 years in the Australian sleepy lizard (*Tiliqua rugosa*) [3]. It has been well established that the longevity of a partnership [4,5], pair coordination, and the compatibility of partners [6] will positively affect measures of reproductive success at the pair level. Whilst such findings have demonstrated that social partnerships can be maintained over long periods of time, the fine scale temporal and spatial associations between partners have received less attention.

The Australian zebra finch (*Taeniopygia castanosis*) is a widely studied bird in captivity, with several studies focusing on the importance of the strength of the pair bond for reproductive success [5–7]. Yet while the zebra finch is the best-studied songbird in captivity, the avian supermodel, little is known about their natural spatial and social organization [8]. Moreover, many characteristics of zebra finches living in the wild contrast starkly with how birds are kept in the laboratory. In the wild, zebra finches are nomadic, moving throughout semi-arid regions in search of grass seed and water [9]. Zebra finches are non-territorial, breed in loose colonies and synchronize their timing of breeding with their neighbours [10]. They are mostly seen alone with their partner but forage in small flocks and use communal, social areas where they gather temporarily in larger groups [9,11,12]. Though both males and females display individually recognizable contact calls [13], these are short range signals (i.e., the loudest call type, the distance call, is only detectable within about 14 m) and so cannot effectively serve to reunite partners once separated in an open landscape [14]. As such, given the natural conditions experienced by zebra finches, it is unclear how pairs maintain partnerships in the wild.

Here, we use automated radio tracking to monitor the movement patterns of wild zebra finch pairs. Our objective was to quantify the degree of spatial cohesion of pairs. To do so, we examined space use and movement patterns between zebra finch pairs compared to neighbouring birds. We predicted that pairs would display higher home range overlap compared to neighbours. Additionally, we predicted that pairs would remain in close proximity throughout the home range.

## II. Methods

Research was approved by the Macquarie University Animal Ethics Committee and covered by a New South Wales DPIE Scientific License SL100378, Ethics ARA 2018/027, and ABBBS Banding Permit.

### a) Data collection

Fieldwork was conducted at Fowlers Gap Research Station, New South Wales, (30°56’58”S, 141°46’02”E) from September to October in 2022 and 2023. During these periods, zebra finches were actively breeding in nest boxes as well as natural nests. In 2022, adult zebra finches were caught by mist nets at three sub-colony sites (each sub-colony being at least 500 m away from another sub-colony) within the larger Gap Hills breeding colony. Each sub-colony site has 30 nest boxes as well as vegetation suitable for natural nests. A female and a male caught simultaneously within a net were considered a pair based on previous observations that zebra finches most commonly are seen in pairs [11,12,15] and that breeding partners are commonly seen flying together [9]. Though not all these ostensible pairs showed the expected patterns (see Results), to avoid circularly defining pairs, however, all male-female pairs caught simultaneously were considered to be mated pairs for all analyses. However, determining pairs via this method might be biased towards catching only those pairs that are especially well-coordinated. To control for this potential bias, in 2023 we caught five additional breeding pairs at two sub-colony sites using nest-box traps during the nestling provisioning period, when nestlings were between six and 11 days old. As a precaution against nest-abandonment and to avoid unnecessary disruption of nestling provisioning, we caught only one provisioning adult per day. After both pair members were radio-tagged, they were confirmed to be visiting together and feeding the nestlings.

Birds were weighed with a 20 g Pesola and, if above 10 g (all adults weighed during this study were >10 g), were tagged with a 0.48 g solar-powered radio tag with a nitinol antenna in 2022 and with a nylon coated braided steel antenna in 2023 (Cellular Tracking Technologies LifeTag, New Jersey, USA). Tags were attached with a stretchable nylon leg-loop harness [16] and were programmed to signal every 5 seconds. Though 12 pairs were tagged in 2022, the antenna of one tag broke after 1 day, which was indicated by a substantial decrease in signal strength and the number of nodes detecting the tag, and so we excluded this individual and the partner from our analyses. In 2022, the remaining tags operated normally for between five to 29 days before a similar change in signal transmission strength was observed indicating the antenna likely broke off.

Immediately after tagging, all individuals were observed to fly away normally. In 2023, when breeding pairs were tracked during chick provisioning, all individuals resumed provisioning shortly after tagging and were observed provisioning for at least five subsequent days, suggesting that there were at least no short-term behavioural impacts of the tags.

### b) Automated radio tracking

To monitor tagged individuals, an automated radio tracking system was installed within the study site, covering ∼1.5 km^2^. 70 radio receivers (Cellular Tracking Technologies Node v2, New Jersey, USA) were mounted every 150 m in a triangular grid with 23 additional receivers placed in areas expected to have increased zebra finch activity, such as near nest boxes, water sources, and areas of grass used for foraging. Receivers recorded the tag ID, the received signal strength (RSS), and the time of the detection. Receiver data was then sent to a central base station.

Tag locations were estimated using RSS-based multilateration. To calibrate the RSS-distance relationship, six tags were held at set distances (1, 2, 5, 10, 15, 25, 50, 75, 100, 150 m) from four receivers for 2 minutes at each distance interval. The common logarithm of RSS was then modelled as a function of distance, yielding the relationship:

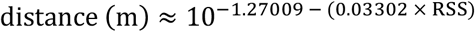

which was used to estimate the Euclidean distance from each receiver. To estimate distance, RSS values were first averaged within a 15 second window with a -5 second lag to remove large fluctuations in signal strength due to interference or multipathing. Within each 15 second interval, the averaged RSS value from receivers within 200 m of the receiver with the strongest averaged signal strength were retained following Paxton et al. [17]. If at least three receivers detected a tag within the interval, then a non-linear least squares model was fit to estimate the location of the tag [17]. For each interval, this process was repeated 100 times by sampling from ± 1 SE around the mean distance for a given RSS-value. Since we calibrated the RSS-distance relationship up to 150 m, distance estimates beyond 150 m were truncated to this value. From the 100 replicated localizations, an error ellipse corresponding to the 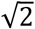-sigma ellipse of a bivariate normal distribution was calculated to estimate the uncertainty around each localization. Overall localization accuracy was assessed through field calibrations which indicated a median difference of 35 m between the estimated and predicted coordinates of tags at known locations.

### c) Data analysis

We removed outlying points that were greater than 2.5 km from the median coordinates of the localizations for each tracked individual. Continuous-time movement models (ctmm) were then fit to these filtered location estimates with their associated error ellipses to identify the autocorrelation structure of the data. The best fitting model was selected based on AIC and used in autocorrelated kernel density estimation (AKDE) to derive home range estimates for each individual, using the R package ‘ctmm’ [18,19]. To assess the home range similarity within and between pairs, we calculated the overlap between home ranges using the Bhattacharyya coefficient, which measures the similarity between two probability distributions [20]. The pairwise estimated home range overlap was compared between partners and neighbouring birds (i.e., other individuals caught within the same sub-colony site) using a mixed-effect beta regression model with individual and partner ID as crossed random effects using the R package ‘glmmTMB’ [21]. This model was then compared to a model excluding group type (i.e., pair or neighbour) as a predictor using a likelihood-ratio test (LRT).

Next, we examined the extent to which pairs moved together throughout their home range to assess spatiotemporal associations between partners. To do so, we calculated a proximity ratio for each possible pair of two individuals (86 dyads in total) using the ‘ctmm’ function *proximity*. The proximity ratio compares the observed separation distances between observed movement paths for dyads to the separation distances from simulated paths assuming random movement. Proximity ratios less than 1 indicate that two individuals are closer together on average than expected given independent movement whereas values greater than 1 indicate avoidance and ratios overlapping 1 indicate independent movement. We fit a linear mixed effect model to examine whether proximity ratios differed between pairs and neighbours with sub-colony site as an additional fixed effect and individual and partner ID as crossed random effects using the R package ‘lme4’ [22]. Observations were weighted by the duration that each dyad was simultaneously tracked. Models without group type (i.e., pair or neighbour) and without sub-colony site were then compared to the full model using a likelihood ratio test. We also examined whether time of day (minutes post-sunrise, log-transformed) influenced the proximity between pairs using a hierarchical generalized additive model [23] with a Gaussian distribution, which we fit with the ‘mgcv’ R package [24]. We specified a global smoother for time as well as factor smoothers where the effects may vary between pairs. This structure is analogous to random slopes in mixed models [23].

## III. Results

The 16 tagged pairs were tracked for an average of 13 days (range 4 to 44 days). During this period, individuals were localized during daylight hours on average every 52 seconds (standard deviation 365 seconds) with longer intervals between detections occurring occasionally when a tag was not detected, likely due to insufficient solar power. During the periods where both partners were located, 29% of each pairs’ localizations were simultaneous, on average. Across groups, the median time between localizations by both partners was eight minutes with 48% of localizations occurring within one minute of the partner.

Overall, across all simultaneous localizations, the median distance between pair members was 56 m compared to 326 m between neighbouring individuals (i.e., individuals caught at the same sub-colony site). There was, however, substantial variation between pairs. For most pairs, the median separation distance between partners was 60 m, but upwards of 600 m in one case (Figure 1).

**Figure 1.**
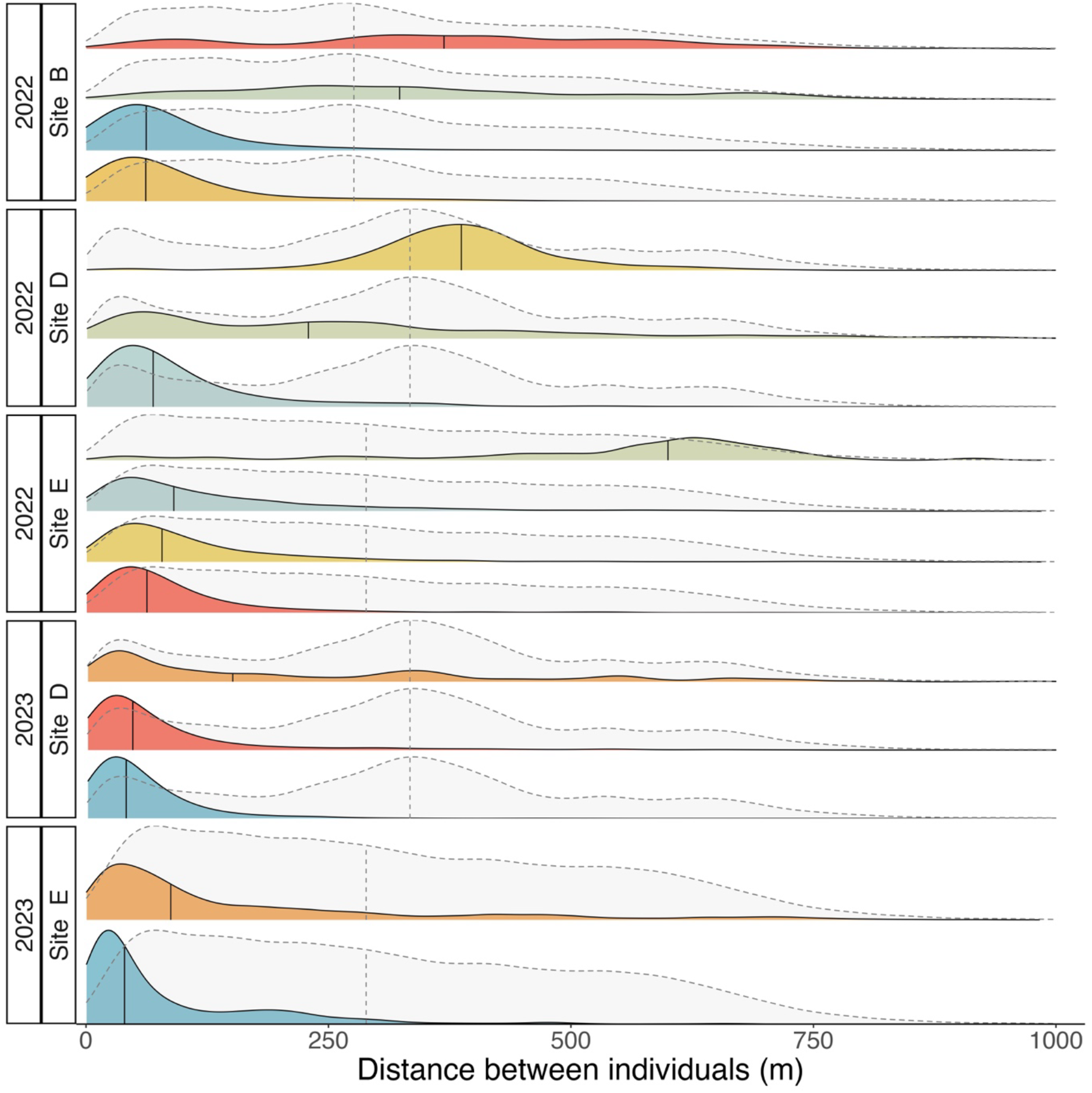
Density plots of the distance (m) between individuals for all simultaneous localizations of pair members (coloured ridges) compared to the distance between neighbours (light grey ridges) at each sub-colony site (B, D or E). Vertical lines indicate the median values for each pair (solid) compared to neighbours in the same section (solid). Each density plot is coloured according to the pair ID, which corresponds to the colour used for each pair in Figures 2 and 3. Pairs were defined as a female and male caught simultaneously in a mist net in 2022 or caught while provisioning nestlings in the same nest-box in 2023.

The average home range area of tracked individuals was 0.45 km^2^ (95% CI: 0.40 -0.49 km^2^, Figure 2A-C). The median home range overlap of pairs was significantly higher for pairs (0.88 Bhattacharyya coefficient) than between neighbouring individuals (0.72 Bhattacharyya coefficient; LRT, p < 0.001; Figure 2D).

**Figure 2.**
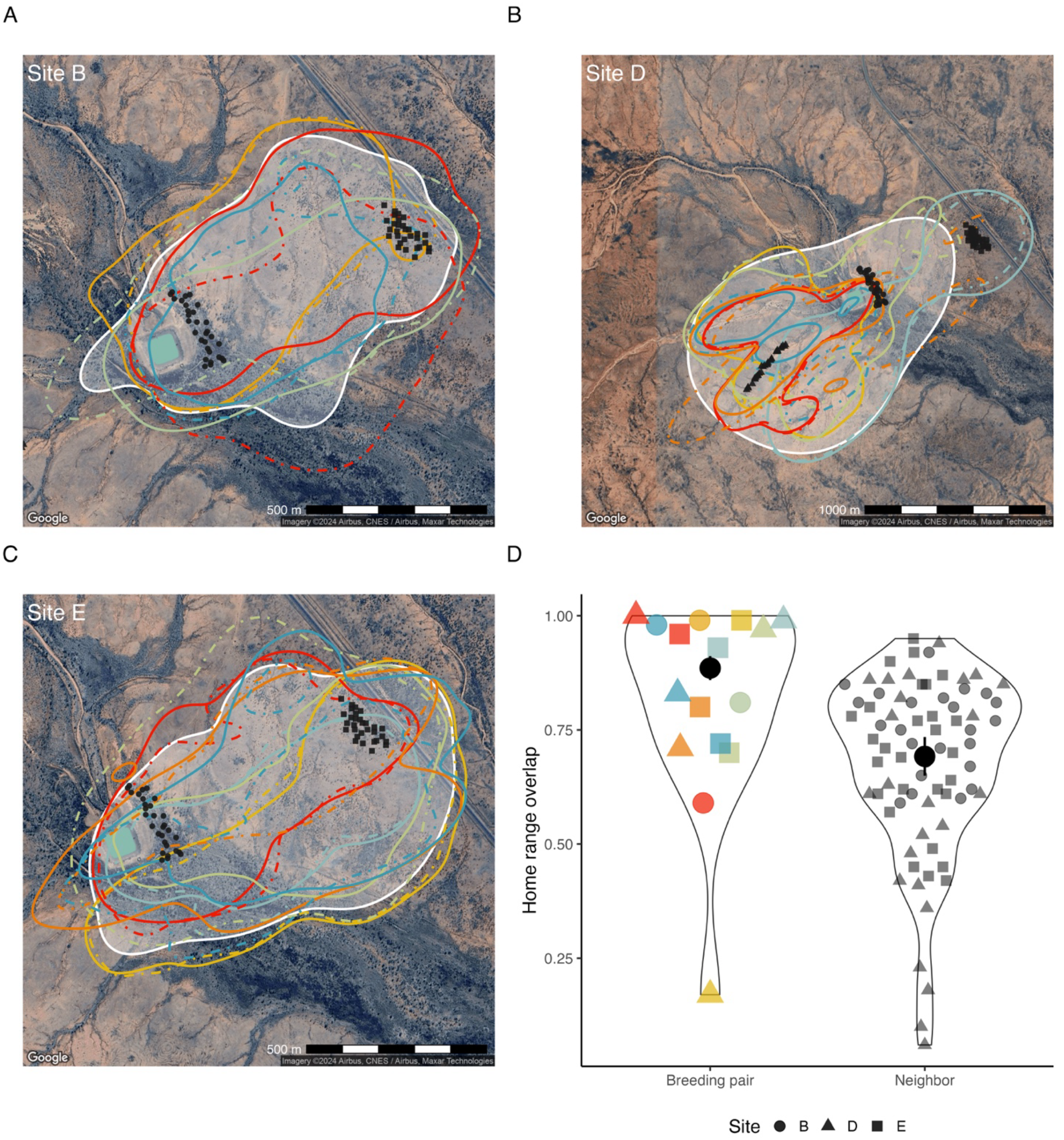
A-C) Home range areas (95% utilization distribution contour) for pairs in each of the three sub-colony sites (B, D, E) in 2022 and 2023. Lines (males, solid; females, dashed) are coloured for each pair. Shaded white regions show the mean population home range for all individuals (as a comparison for neighbours) in each sub-colony site. Black points show the nest boxes within each sub-colony site with shapes corresponding to the sub-colony site (legend shown in 2D). Satellite imagery was obtained using the R package ‘ggmaps’ [25]. D) Violin plots of the home range overlap (of pairs and neighbouring pairs) from a mixed-effect beta regression model of home range overlap as a function of each pair group. Coloured points (pairs) and grey points (neighbours) show the raw estimated home range overlap for each pair, black points show the mean ± SE. For pairs, the shape corresponds to the sub-colony site where it was caught.

The majority of dyads (82 out of the 86 possible) had estimated proximity ratios less than 1, indicating that birds were closer on average than expected given independent movement (Figure 3A). Moreover, pairs had significantly lower proximity ratios compared to neighbouring birds (LRT, p < 0.001). A hierarchical generalized additive model indicated that separation distance was not significantly affected by time of day (p = 0.229, Figure 3B).

**Figure 3.**
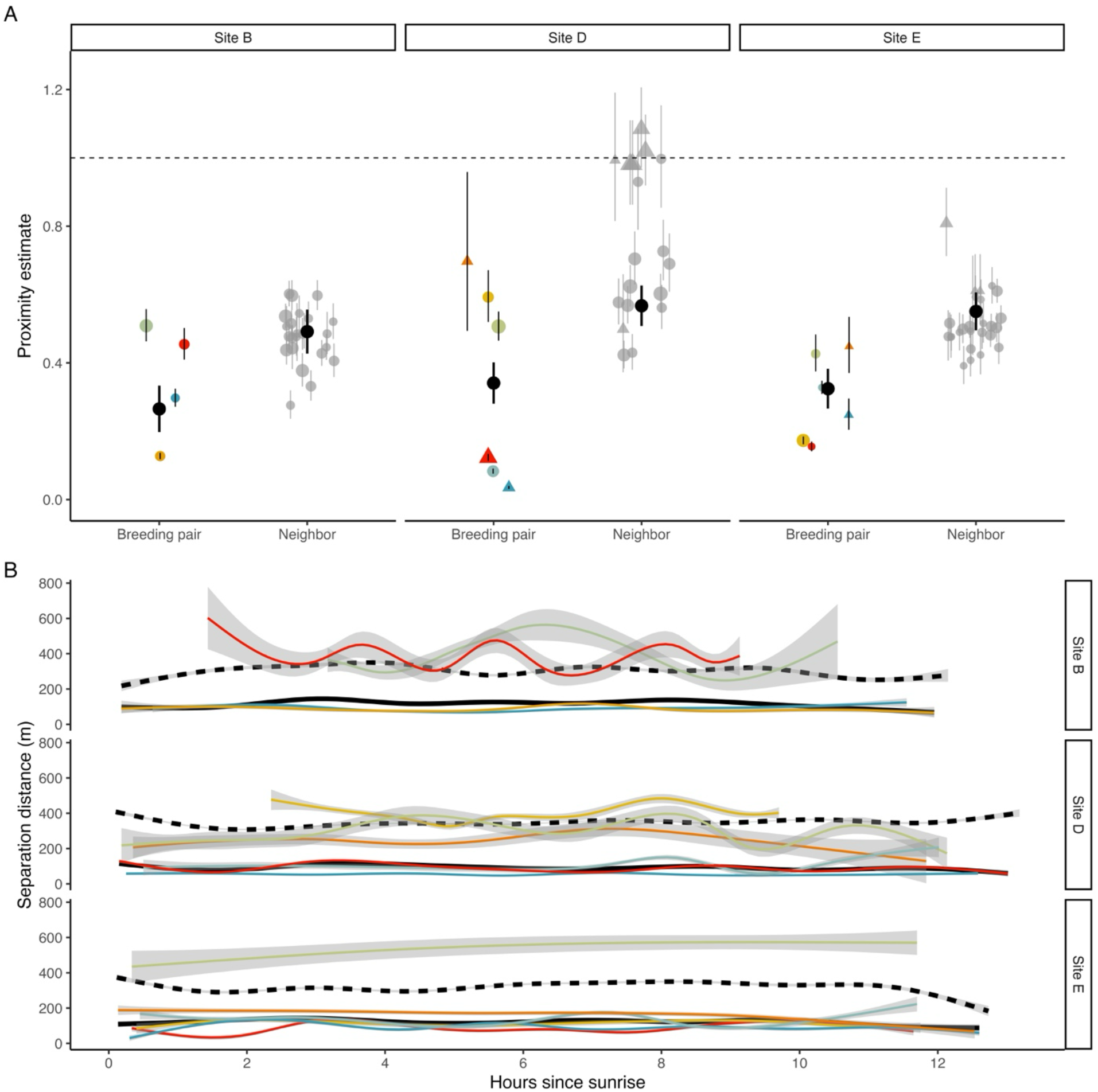
A) Proximity estimates for pairs and neighbours in the three sub-colony sites. Black points show the modelled mean (with 95% CI) proximity estimate for each group. Pairwise proximity estimates (with 95% CI) for pairs (coloured points, corresponding to pair ID) and neighbours (grey points) are also shown. Dyads from 2022 are shown as circles and dyads from 2023 are shown as triangles. Points are scaled according to the number of days each dyad was simultaneously tracked (1.5 to 28 days). Proximity estimates less than 1 correspond to individuals being closer on average than expected given independent movement whereas values greater than 1 indicate that individuals are farther apart than expected. B) Observed separation distances for pairs as a function of hours since sunrise. Coloured lines show pairs (with 95% CI) and black lines (pairs, solid; neighbours, dashed) show the modelled group mean values.

## IV. Discussion

Using automated radio tracking to characterize the movement patterns of zebra finch pairs in the wild, we observed that multiple measures of space-use were substantially more similar for pairs than between neighbouring individuals. Home range overlap was significantly greater for pairs compared to neighbouring individuals, though pairs were clearly not territorial and home range overlap with neighbouring birds was high. Beyond sharing similar home ranges, pairs also remained within close proximity throughout the home range, significantly more so than compared with neighbouring birds. This proximity was maintained throughout the day for paired birds. As such, despite conservatively classifying some pairs based on a single instance where a male and a female were simultaneously captured, which may have misclassified 3 pairs (out of 11) in one year of the study, we nevertheless observed that partners displayed significantly stronger spatial cohesion compared to neighbouring individuals. Taken together, these findings indicate that in the wild, zebra finch pairs maintain high levels of spatial cohesion across large areas during the breeding season.

Unlike many passerine species, zebra finches are non-territorial during the breeding period [9,26]. This is consistent with our finding that overall, home range overlap was high for neighbouring birds (mean overlap of 72%). This lack of territoriality is likely related to their multi-level social structure [14] and the distribution of locally abundant water and grass seeds within the arid and semi-arid regions inhabited by zebra finches, which does not permit (or warrant) pairs to exclude competitors [9]. Water points are patchily distributed at isolated sources, such as naturally occurring ephemeral ponds, and also at anthropogenic sources, such as livestock troughs, tanks, and dams (as seen in Figure 2A-C); none of which could be defended as part of a territory by zebra finches. Additionally, as obligate granivores, zebra finches forage in open grasslands where seed is relatively homogeneously distributed over large areas. Such ecological factors presumably account for the relatively large mean home range zebra finches (0.47 km^2^), compared to home ranges for generalist species breeding in temperate regions, such as great tits (*Parus major*, 0.02 km^2^) where resources can be locally defended [27]. Furthermore, despite sharing largely similar home ranges, within this area pairs maintained significantly closer proximity. As such, this was not due to pairs remaining on a defined territory, but rather because pairs used this relatively large home range area together in a coordinated way. Still, though pair proximity was greater compared to neighbours, proximity estimates between neighbouring birds indicated that neighbours were closer together than would be expected given independent movement, which suggests that neighbouring birds concurrently visited similar areas, such as foraging and drinking locations or social hangout sites [11]. Instead of competing for these locally abundant resources, it is likely that birds benefit from exploiting them in the safety of a group, which is reflected in the home range overlap and movements of both pairs and neighbouring individuals.

Continually maintaining close proximity to a partner may serve multiple functions in zebra finches. Pairs that constantly maintain reproductive readiness would be able to rapidly respond to unpredictably occurring breeding conditions and so could maximize the number of clutches within a breeding season [7]. In captivity, experimentally isolating zebra finch partners increases plasma corticosterone levels, which only returns to pre-separation baseline levels after partners have been reunited for 48 hours [28]. Given that corticosterone tends to inhibit breeding and parental care behaviours, maintaining pair cohesion is likely critical to facilitating rapid reproduction. Additionally, as wild zebra finch pairs are nomadic and might undertake extensive movements to locate water sources [9], continual contact with a partner would facilitate pairs jointly relocating. During breeding, spatial cohesion may have multiple benefits as well, such as augmenting nest visit synchrony, which has been linked with higher reproductive success in zebra finches [15], or potentially by increasing predator awareness while foraging with a pair member [29]. Taken together, these possible benefits suggest that maintaining constant proximity is a critical behaviour of wild zebra finch pairs. The degree of spatial cohesion, however, will likely vary throughout the breeding season. For instance, while spatial cohesion may be high during pre-breeding in order to maintain breeding readiness, during incubation it will likely decrease as partners trade-off incubation stints [9], and during provisioning there might be more variation between pairs. In this study, we could not fully explore the sources of this variation as the breeding status of most pairs was unknown (in 2022) or did not vary (all pairs in 2023 were provisioning nestlings). We did observe though that pairs tracked during provisioning showed similar patterns in terms of spatial proximity and home range overlap compared to birds tracked during an unknown breeding stage. Still, given that spatial cohesion may serve various functions, quantifying spatial cohesion at different stages of the annual cycle would be a valuable next step in identifying the adaptive benefits of this behaviour and its role in maintaining continuous partnerships.

Although sexual conflict received significant attention in the behavioural ecological research on socially monogamous mating systems during the previous decades, there is growing appreciation for the importance of cooperation between breeding partners in many species [30]. In socially monogamous species with long-term partnerships and low rates of extra-pair paternity [31], such as the zebra finch, partners will share a greater proportion of each other’s lifetime reproductive output. In this situation, the evolutionary interests of pairs are more aligned, which will favour cooperation and coordination between pair members [30]. Spatial cohesion is a behavioural interaction where both partners must coordinate to remain in contact. To date, only one other study (to our knowledge) has simultaneously tracked the fine-scale spatial movements of songbird pairs, where it was shown that great tit pairs coordinate provisioning by utilizing the same core area around the nest during foraging [32]. In great tits, however, spatial cohesion is suggested to enable partners to monitor one another’s provisioning levels and thereby to reduce sexual conflict over parental care. Additionally, great tits are more likely to alternate provisioning bouts, in a tit-for-tat fashion, rather than synchronously provisioning as zebra finches do [15,33]. Nonetheless, given that in both of the two studies tracking fine-scale movements of songbird pairs it was found that pairs show high spatio-temporal synchrony, this suggests that such behaviours may be widespread. These findings demonstrate that spatial cohesion may play an important role in many socially monogamous partnerships.

## Acknowledgments

We are grateful to Elke Molenaar, Emma Scheltens, Evelien ter Avest, Lyanne Brouwers, Riccardo Ton, and Rita Fragueira for fieldwork assistance. We also thank Garry Dowling and Mark Tilley for assistance installing components of the automated radio tracking system and Vicki Dowling for logistical support. The project was supported in part by a grant by the Dutch Research Council (NWO) to MN (ALWOP.334).

## References

1. Van De Pol M, Heg D, Bruinzeel LW, Kuijper B, Verhulst S. 2006 Experimental evidence for a causal effect of pair-bond duration on reproductive performance in oystercatchers (Haematopus ostralegus). Behav. Ecol. 17, 982–991. (doi:10.1093/beheco/arl036)

2. Jeschke JM, Wanless S, Harris MP, Kokko H. 2007 How partnerships end in guillemots Uria aalge: Chance events, adaptive change, or forced divorce? Behav. Ecol. 18, 460–466. (doi:10.1093/beheco/arl109)

3. Leu ST, Burzacott D, Whiting MJ, Bull CM. 2015 Mate familiarity affects pairing behaviour in a long-term monogamous lizard: Evidence from detailed bio-Logging and a 31-year field study. Ethology 121, 760–768. (doi:10.1111/eth.12390)

4. Sánchez-Macouzet O, Rodríguez C, Drummond H. 2014 Better stay together: Pair bond duration increases individual fitness independent of age-related variation. Proc. R. Soc. B Biol. Sci. 281. (doi:10.1098/rspb.2013.2843)

5. Adkins-Regan E, Tomaszycki M. 2007 Monogamy on the fast track. Biol. Lett. 3, 617–619. (doi:10.1098/rsbl.2007.0388)

6. Schuett W, Dall SRX, Royle NJ. 2011 Pairs of zebra finches with similar ‘personalities’ make better parents. Anim. Behav. 81, 609–618. (doi:10.1016/j.anbehav.2010.12.006)

7. Maldonado-Chaparro AA, Forstmeier W, Farine DR. 2021 Relationship quality underpins pair bond formation and subsequent reproductive performance. Anim. Behav. 182, 43–58. (doi:10.1016/j.anbehav.2021.09.009)

8. Griffith SC, Buchanan KL. 2010 The Zebra Finch: The ultimate Australian supermodel. Emu. 110. (doi:10.1071/MUv110n3_ED)

9. Zann RA. 1996 The zebra finch: a synthesis of field and laboratory studies. Oxford: Oxford University Press.

10. Brandl HB, Griffith SC, Schuett W. 2019 Wild zebra finches choose neighbours for synchronized breeding. Anim. Behav. 151, 21–28. (doi:10.1016/j.anbehav.2019.03.002)

11. Loning H, Fragueira R, Naguib M, Griffith SC. 2023 Hanging out in the outback: the use of social hotspots by wild zebra finches. J. Avian Biol. 2023. (doi:10.1111/jav.03140)

12. McCowan LSC, Mariette MM, Griffith SC. 2015 The size and composition of social groups in the wild zebra finch. Emu 115, 191–198. (doi:10.1071/MU14059)

13. Elie JE, Theunissen FE. 2018 Zebra finches identify individuals using vocal signatures unique to each call type. Nat. Commun. 9. (doi:10.1038/s41467-018-06394-9)

14. Loning H, Griffith SC, Naguib M. 2024 The ecology of zebra finch song and its implications for vocal communication in multi-level societies. Philos. Trans. R. Soc. B Biol. Sci. 379. (doi:10.1098/rstb.2023.0191)

15. Mariette MM, Griffith SC. 2012 Nest visit synchrony is high and correlates with reproductive success in the wild zebra finch Taeniopygia guttata. J. Avian Biol. 43, 131–140. (doi:10.1111/j.1600-048X.2012.05555.x)

16. Bircher N, Van Oers K, Hinde CA, Naguib M. 2020 Extraterritorial forays by great tits are associated with dawn song in unexpected ways. Behav. Ecol. 31, 873–883. (doi:10.1093/BEHECO/ARAA040)

17. Paxton KL, Baker KM, Crytser ZB, Guinto RMP, Brinck KW, Rogers HS, Paxton EH. 2022 Optimizing trilateration estimates for tracking fine-scale movement of wildlife using automated radio telemetry networks. Ecol. Evol. 12. (doi:10.1002/ece3.8561)

18. Calabrese JM, Fleming CH, Gurarie E. 2016 ctmm: an <scp>r</scp> package for analyzing animal relocation data as a continuous-time stochastic process. Methods Ecol. Evol. 7, 1124–1132. (doi:10.1111/2041-210X.12559)

19. Fleming CH, Calabrese JM. 2017 A new kernel density estimator for accurate home-range and species-range area estimation. Methods Ecol. Evol. 8, 571–579. (doi:10.1111/2041-210X.12673)

20. Winner K, Noonan MJ, Fleming CH, Olson KA, Mueller T, Sheldon D, Calabrese JM. 2018 Statistical inference for home range overlap. Methods Ecol. Evol. 9, 1679–1691. (doi:10.1111/2041-210X.13027)

21. Brooks ME, Kristensen K, van Benthem KJ, Magnusson A, Berg CW, Nielsen A, Skaug HJ, Mächler M, Bolker BM. 2017 glmmTMB balances speed and flexibility among packages for zero-inflated generalized linear mixed modeling. R J. 9, 378–400. (doi:10.32614/rj-2017-066)

22. Bates D, Maechler M, Bolker B. 2012 lme4: Linear mixed-effects models using S4 classes.

23. Pedersen EJ, Miller DL, Simpson GL, Ross N. 2019 Hierarchical generalized additive models in ecology: An introduction with mgcv. PeerJ 2019. (doi:10.7717/peerj.6876)

24. Wood SN. 2011 Fast stable restricted maximum likelihood and marginal likelihood estimation of semiparametric generalized linear models. J. R. Stat. Soc. Ser. B Stat. Methodol. 73, 3–36. (doi:10.1111/j.1467-9868.2010.00749.x)

25. Kahle D, Wickham H. 2013 ggmap: Spatial visualization with ggplot2. R J. 5, 144–161. (doi:10.32614/rj-2013-014)

26. Tobias JA, Sheard C, Seddon N, Meade A, Cotton AJ, Nakagawa S. 2016 Territoriality, social bonds, and the evolution of communal signaling in birds. Front. Ecol. Evol. 4, 1–15. (doi:10.3389/fevo.2016.00074)

27. Naguib M, Titulaer M, Waas JR, van Oers K, Sprau P, Snijders L. 2022 Prior territorial responses and home range size predict territory defense in radio-tagged great tits. Behav. Ecol. Sociobiol. 76. (doi:10.1007/s00265-022-03143-3)

28. Remage-Healey L, Adkins-Regan E, Romero LM. 2003 Behavioral and adrenocortical responses to mate separation and reunion in the zebra finch. Horm. Behav. 43, 108–114. (doi:10.1016/S0018-506X(02)00012-0)

29. Beauchamp G. 2002 Higher-level evolution of intraspecific flock-feeding in birds. Behav. Ecol. Sociobiol. 51, 480–487. (doi:10.1007/s00265-002-0461-7)

30. Griffith SC. 2019 Cooperation and coordination in socially monogamous birds: moving away from a focus on sexual conflict. Front. Ecol. Evol. 7, 1–15. (doi:10.3389/fevo.2019.00455)

31. Griffith SC, Holleley CE, Mariette MM, Pryke SR, Svedin N. 2010 Low level of extrapair parentage in wild zebra finches. Anim. Behav. 79, 261–264. (doi:10.1016/j.anbehav.2009.11.031)

32. Baldan D, van Loon EE. 2022 Songbird parents coordinate offspring provisioning at fine spatio-temporal scales. J. Anim. Ecol. (doi:10.1111/1365-2656.13702)

33. Mariette MM, Griffith SC. 2015 The adaptive significance of provisioning and foraging coordination between breeding partners. Am. Nat. 185, 270–280. (doi:10.1086/679441)

